# Close range vocal interaction in the common marmoset (*Callithrix Jacchus*)

**DOI:** 10.1101/2019.12.19.882118

**Authors:** R. Landman, J. Sharma, J.B. Hyman, A. Fanucci-Kiss, O. Meisner, S. Parmar, G. Feng, R. Desimone

## Abstract

Vocal communication in animals often involves taking turns vocalizing. In humans, turn taking is a fundamental rule in conversation. Among non-human primates, the common marmoset is known to engage in antiphonal calling using phee calls and trill calls. Calls of the trill type are the most common, yet difficult to study, because they are not very loud and uttered in conditions when animals are in close proximity to one another. Here we recorded trill calls in captive pair-housed marmosets using wearable microphones, while the animals were together with their partner or separated, but within trill call range. Trills were exchanged mainly with the partner and not with other animals in the room. Animals placed outside the home cage increased their trill call rate and uttered more trills in response more to their partner. The fundamental frequency, F0, of trills increased when animals were placed outside the cage. Our results indicate that trill calls can be monitored using wearable audio equipment. Relatively minor changes in social context affect trill call interactions and spectral properties of trill calls, indicating that marmosets can communicate subtle information to their partner vocally.

## Introduction

Turn-taking is a fundamental feature of human conversation (1). Similar temporal regulation of vocal interactions can also be observed in non-human primates (2,3) and birds (4). In the Common Marmoset (Callithrix Jacchus), turn-taking is observed in phee and trill calls (2,5–7). Trill calls are relatively quiet calls, uttered in conditions when animals are in close proximity to one another, making them more difficult to attribute to the caller than phee calls. Although trill calls are thought to have a function in marmoset communication, it is been unclear whether trill calls change in response to subtle changes in the social group. Here we have used wearable voice recorders to allow detection and attribution of trill calls to the specific animals that uttered them.

Marmosets are a small bodied, highly social and vocal species of New World monkey native to South America. In the wild, they live in social groups of up to 15 individuals, usually consisting of one breeding pair and up to several generations of offspring. These primates have a cooperative breeding system in which individuals other than the mother help caring for infants (8,9). Mainly due to their arboreal habitat, these primates communication is via vocalization for which they use a rich call repertoire consisting of approximately 13 different call types (10). The *phee* and the trill are thought of as contact calls, serving to monitor the presence of group members. The *phee* call is evoked when an animal is far removed from other marmosets and not in visual contact (2,6). The trill call occurs often when animals are in close proximity from one another (6,7,11). Antiphonal calling, which means exchanging calls between individuals back and forth, happens with both phee calls and trill calls (6,7,11).

Paradigms in which antiphonal phee calling is evoked have proven useful in elucidating mechanisms of social interaction, vocal development and the evolution of language (12,13). Phee calls and phee interactions are learned from parents (13), suggesting that this could be a good model of human conversational turn taking. Evidence suggests that marmosets can also recognize one another by the sound of their phee calls (14). Phee call rate is affected by changes in social context (15): While phees were uncommon when animals were in their native group (in captivity), phee rate was elevated for several weeks after being paired with a new animal. Short term (10min) isolation is also known to increase phee rate (5,13,15).

Less is known about trill calls than about phee calls, particularly whether they are affected by social context. Trills from one animal are often followed by trills from other animals at a latency of less than 1 second (6). In a study with three pygmy marmosets, Snowdon & Cleveland (16) found that within trill call bouts, certain sequences were more common than others, suggesting a rule system that governs the order of antiphonal trill calls. However, is it not clear how common trill interactions are and what role circumstances play.

Audio features of vocalizations often change with arousal, emotion, and distance from peers. The ability to adjust acoustic properties, such as amplitude and frequency, in response to emotions and changes in the environment can be important for communication (17,18). In marmosets, phee calls have been shown to increase in frequency and amplitude with increasing levels of isolation from the group (15). However, it is not known whether any audio features of trill calls are also affected by changes in the environment or social context.

In the present study, we analyzed the temporal relationships and audio features of calls among pair-housed marmosets. We recorded their natural trill call exchanges when together in the home cage and when one animal was in the home cage and the other was in a separate cage about 0.5m away while the animal maintained visual and auditory access to their partner Indeed, we find that this subtle manipulation increases trill call rate and trill call interactions. The fundamental frequency (F0) of the calls increased when animals were separated.

The entire dataset for this study (raw audio + annotations) is downloadable at OSF (https://osf.io/pswqd/, DOI: 10.17605/OSF.IO/PSWQD)

## Methods

### Animals

Twenty marmosets, ranging from 1.5 to 11 years old, were used as subjects. The animals lived in pairs. Five pairs were mixed sex and five pairs were same sex (male siblings). All pairs were housed in a room with other marmoset cages (between 12 and 25 Marmosets in total). A typical room layout is shown in Figure 1A. Home cage size was 77.5 x 77.5 x 147 cm. There was >90 cm of space between neighboring cages (edge to edge). The cages have opaque panels on the side, which somewhat reduce the intensity of sound coming from animals in other cages. There was >250 cm between the focal cage and cages across from them. In separation conditions during the experiment, one animal at a time was placed in a transport cage, size 30 x 30 x 33 cm, with 0-50 cm of space between transport cage and home cage. The transport cage is made of wire mesh with a transparent polycarbonate door. At the time of experimentation all pairs had lived together for at least 6 months. The males in the mixed sex pairs were vasectomized. The animals had ad libitum access to water and received standard daily diet and nutritional supplements. All experimental manipulations were made under institutional guidelines and approved by the MITs Committee for animal care (CAC), the Broad Institute’s Institutional Animal Care and Use Committee (IACUC) and in accordance with the NIH Guide for the Care and Use of Laboratory Animals.

**Figure 1.**
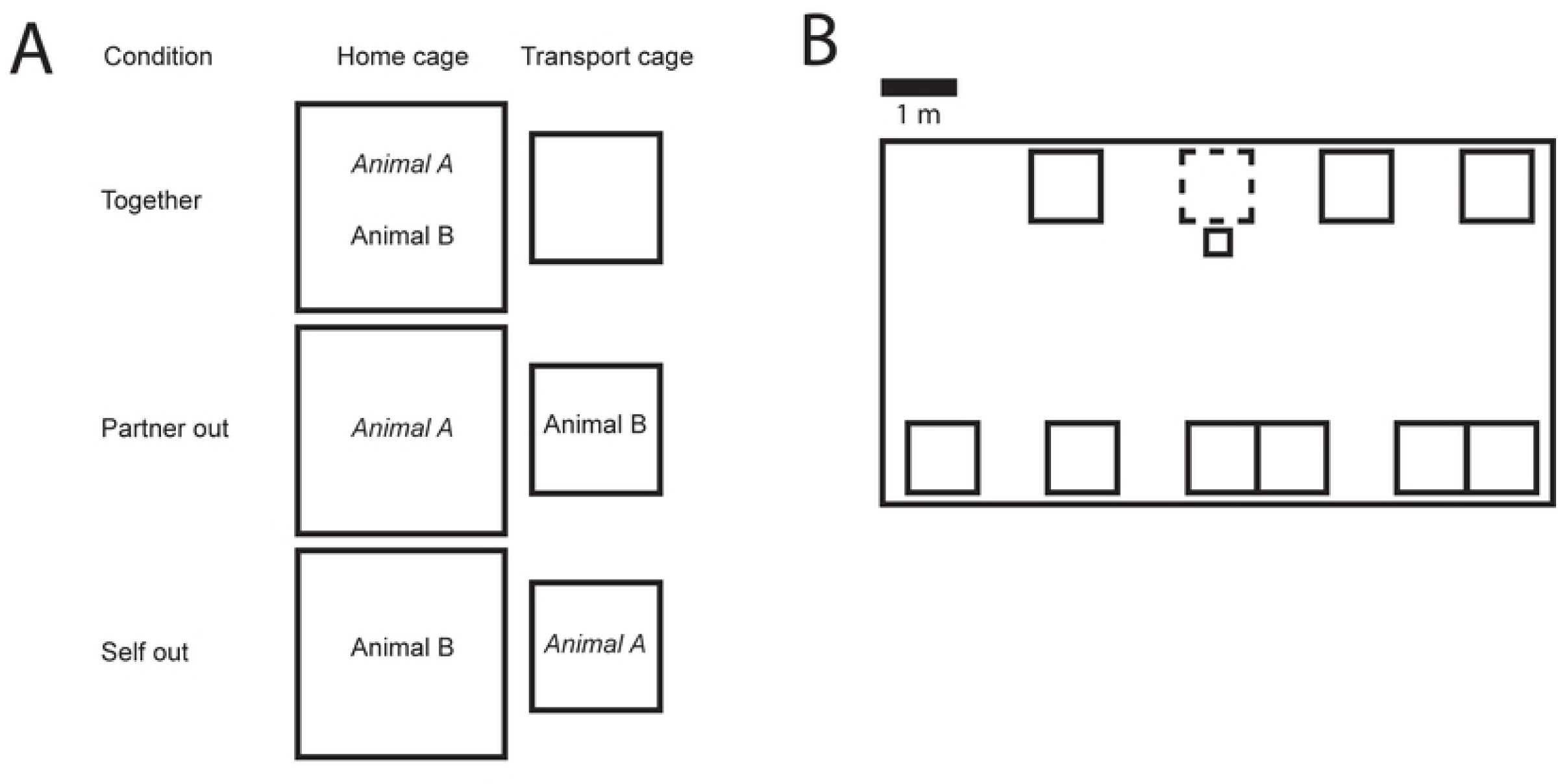
**A**. Illustration of the experimental conditions with Animal A as the focal animal. **B**. A cartoon example of a typical room layout and positioning of the cages. The large rectangle is the room and the small rectangles represent cages. Each single cage has two marmosets. For families, two single cages are connected to make one large cage. The cage with dashed line is the cage recorded from. In front of it is the transport cage where the animal outside of the home cage was.

### Materials

Vocalizations were recorded using commercially available, light weight Panictech and Polend digital voice recorders, mounted using duct tape on a custom made leather jacket. The voice recorder dimensions are 45 x 17 x 5mm and weight is 6.9gr. This device has an omnidirectional electret microphone and samples at 44.1kHz.

### Habituation procedure

Prior to recording, each animal was habituated to wearing a jacket. First, each animal was trained to enter a transport cage for a food reward. The animal was taken out of the transport cage and restrained by one experimenter while another experimenter put on the jacket. All animals were habituated by gradually increasing the duration the jacket was left on from 10 min to approximately 1 hour over a series of 5 or more sessions.

### Recording procedure

Recordings were done with one pair at a time. Each session, a series of 4 epochs was recorded. The first epoch, the animals were together in the home cage. The second epoch, one animal was placed in a transport cage placed on a table 0-30 cm from the home cage. The dyads could see and hear each other during this time. The third epoch, the animals were back together in the home cage, and the fourth epoch, the other animal was placed in the transport cage. Thus, for each animal, we had a condition in which the animal was together with their partner (‘Together’), a condition in which their partner was out (‘Partner out’), and a condition in which they themselves were out (‘Self out’). See Figure 1A for an illustration of the conditions. The order in which animals went out was randomized across sessions. The mean number of sessions per pair was 3.5. Mean epoch duration was 861 seconds.

### Analysis

Wave data from each recorder were uploaded to a computer for analysis. Wave files were manually aligned type in Audacity ® (v. 2.1.0) software. The data was partially annotated by hand, and partially annotated using an auto-detection algorithm (20). Annotations included call start time, call end time, call type, and caller ID (animal A, animal B, or other). A known issue with handheld recorders is inter-device time drift due to errors in oscillators (21). In our recorders the drift was up to 240 ms per hour. This was corrected in Audacity using the ‘change tempo’ effect.

Eight call types were distinguished: *trill, phee, trillphee, chirp, twitter, tsik, ek, and chatter* (10,22). A 9^th^ category named *other* was for calls that did not fit into any category. For call types that often occur in multiple utterances less than 0.5 seconds apart, such as phee, chatter, chirp and twitter, a bout was labeled as a single call. Calls were attributed to animal A, animal B, or to animals from other cages based on 2 considerations: (1) Wave amplitude and spectral intensity. Amplitude and spectral intensity are largest on microphone of the animal that vocalizes. If the amplitude or intensity is low and about the same level on both microphones, the call is likely coming from an animal in a different cage. (2) Sharpness of the spectrogram image. The spectrogram of distant calls appears smeared compared to calls from animals wearing the microphone. Due to the poorer definition and low amplitude of calls attributed to other animals in the room, we were not able to determine call type of those calls. Most difficult to attribute were loud calls, such as phee, because they were often equal in amplitude on both recorders or the signal clipped.

To illustrate the relation between audio signal strength and distance, we show spectrograms from Audacity, the software in which we did the annotation, of a pre-recorded trill played on a speaker at various distances (Figure 2A). Between 0.05 meters (the approximate distance from recorder to mouth of the animal) and 1 meter, there is steep fall off in amplitude and spectral energy. To illustrate this in the context of animals wearing the recorders, in Figure 2B we show a single video frame taken at the moment a trill call was uttered. The video was taken with a 3D camera (ZED mini, Stereolabs Inc.). Audio and video was synced offline by aligning an audio and visual signal played on a tablet computer within the video frame. We calculated real world coordinates of the animals using the ZED SDK and determined that inter-animal distance (distance between the pin and green colored patches) was 549mm. Based on comparison of the spectrograms between the two recorders (Figure 2C), we attributed this call to animal 1 (pink jacket). Another example is shown in Figure 2D-E. Here inter-animal distance is 76mm. Based on comparison of the spectrograms, this call was attributed to animal 2 (green jacket).

**Figure 2.**
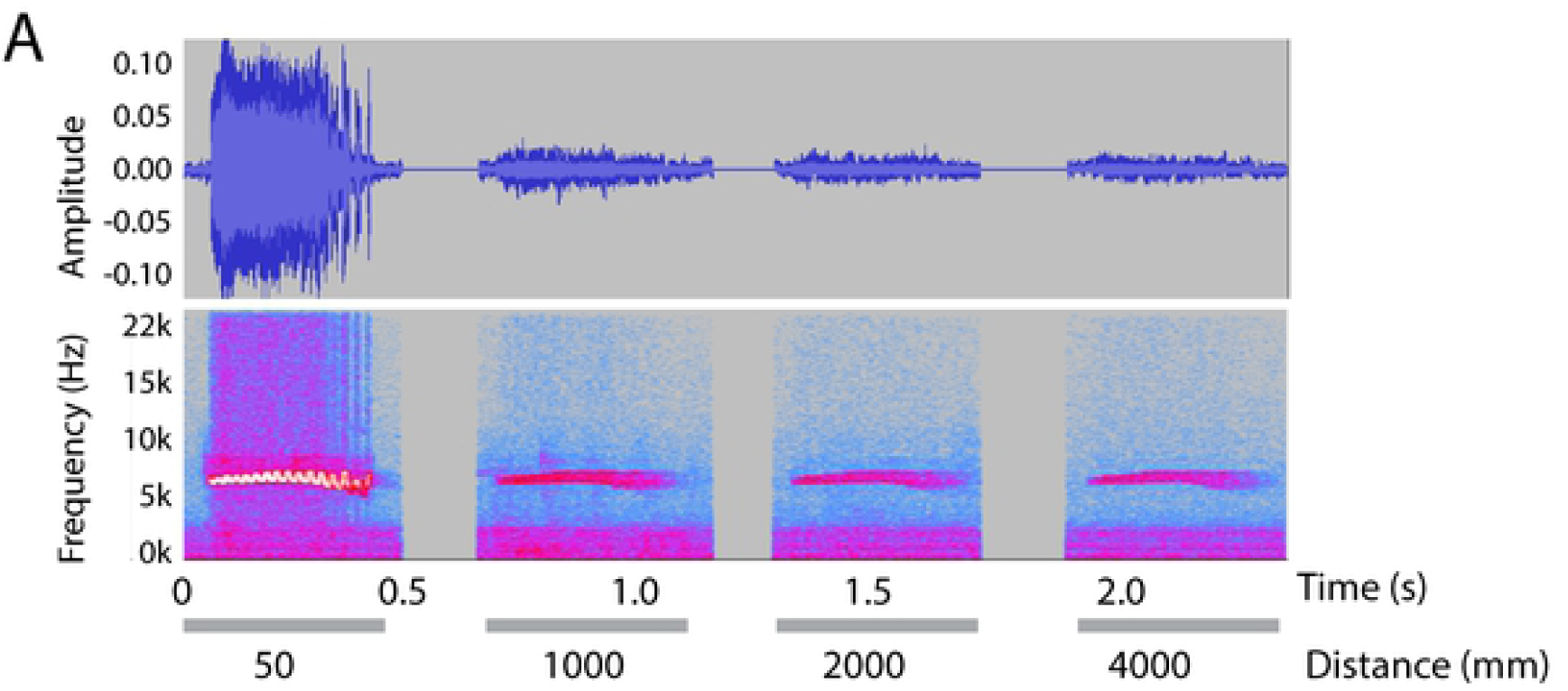
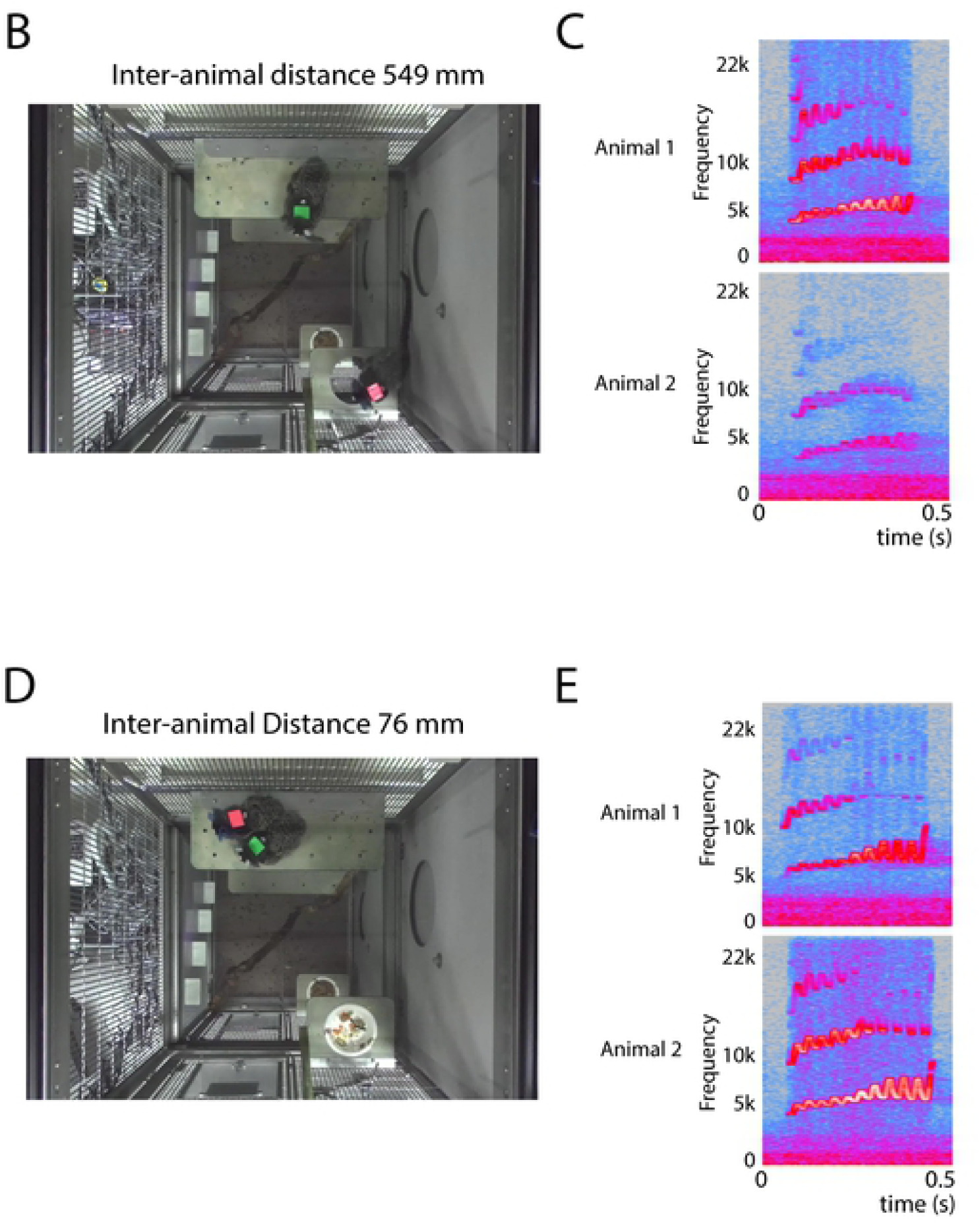
Audio signal strength versus distance. **A**. Wave (upper row) and spectral intensity (bottom row) from one voice recorder as seen in Audacity, the software we used for annotation. Shown is a recording of a single trill call played on a speaker at various distances. There is a steep fall off between 005 m (the approximate distance between the animals’ mouth and the recorder) and 1 m. Beyond 1 m, fall off is less steep. Besides lower intensity, the spectrogram also shows less detail, such as the sinusoidal frequency modulation that is typical of trill calls. **B**. A video frame taken at the onset time of a trill call showing two animals wearing voice recorders marked pink and green. The inter-animal distance is 549 mm. **C**. Spectrograms of the call from the two voice recorders show a difference in intensity that can be used to infer which animal called. This call is attributed to animal 1 (green). D. A video frame with inter-animal distance 76 mm. **D**. Spectrograms show higher intensity for animal 2 (pink), to whom the call is attributed.

As an additional control for source separation between the animals wearing the voice recorders and other animals in the room, a subset of recordings was done with 3-6 additional microphones spread throughout the room, including one aimed at the cage under study. Cross-correlation on rectified audio signals between all possible combinations of microphones was used to determine the microphone with the earliest onset for each call. The location of the cross-correlation peak relative to zero lag was used as the indicator of which microphone had the earlier onset. If the earliest onset came from the microphone aimed at the cage under study, the call was marked as coming from that cage. If the earliest onset came from any other microphone, the call was marked as coming from other animals in the room. Attributions from the microphone array were compared to attributions from the voice recorders.

Call rates were calculated using call onset times binned in 10ms bins. For each animal we calculated mean call rate in 5 s segments following calls from the partner (T=0). The maximum in the period 0 ≤ T ≤ 1 s was taken as the peak. The time bin of the maximum was taken as the peak latency. The peak was considered significant if it was higher than baseline plus two standard deviations, where baseline is call rate across the entire session. Spectrograms of trill calls were made using the *spectrogram* function in Matlab after zero padding to 30000 samples, using a Hamming window of length 256 samples with 128 samples overlap and an FFT length of 1024. Fundamental frequency F0 was calculated based on the call spectra of the initial 150ms of the calls to ensure inclusion of short calls as well as long ones. Spectral peaks were detected using the *findpeaks* function in Matlab, with a *MinPeakProminence* of 0.2 and *MinPeakDistance* of 20. The first (lowest) peak was taken as the fundamental frequency.

Hypothesis testing was done in SPSS, version 25.

## Results

We recorded an average of 850 calls per animal. Calls of the type trill were the most common with a rate of 1.8 calls/minute when animals were together in the home cage (Figure 3). When the animals were separated (i.e. when either member of the pair was in a small cage in front of the home cage), trill call rate increased. A one-way repeated measures ANOVA on trill call rate yielded a significant effect of separation condition, whereas other call types did not (Bonferroni corrected α=0.0045, F(2,38)=7.056, p=0.002). The strongest increase in trill rate was when the animals themselves were outside the home cage (‘self out’).

**Figure 3.**
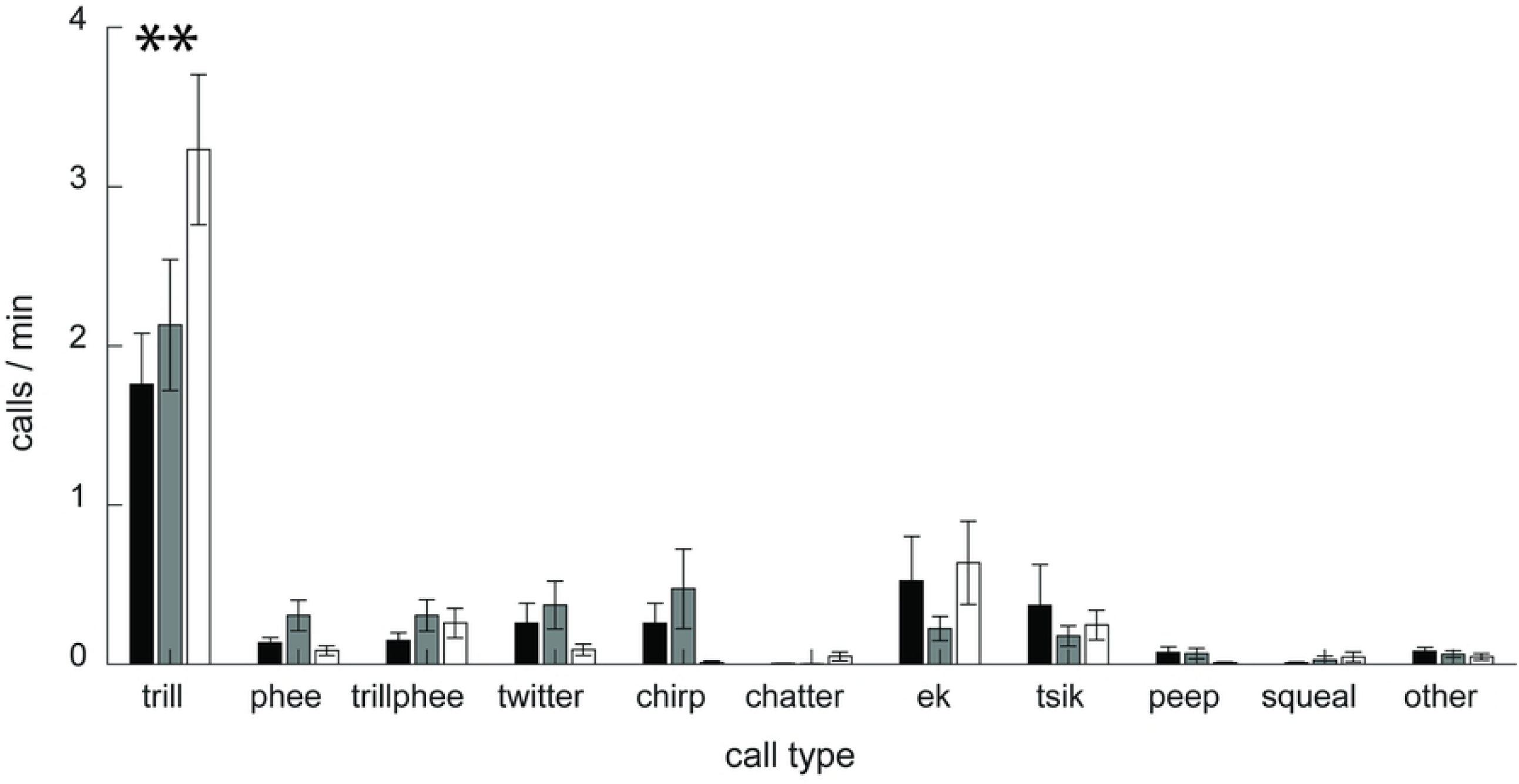
Vocalization rate. Plotted for different call types when animals are together in the home cage (black) or separated, i.e. when the partner is out (gray) or when they themselves are out (white). Trill calls were the most common call type. The rate of trill increased with separation.

By analyzing the relative timing of calls from different individuals, we can determine whether there may be vocal interaction between them. We distinguish between calls from individuals making up each pair and animals in other cages, which could be heard in the background on each of the wearable recorders. Figure 4A shows that call rate contingent upon calls from the partner rises steeply, reaching a peak within about 1 s. Call rate from either partner contingent upon calls from other animals does not show a peak. This indicates that there is a relationship in the timing of vocalizations between cage partners but not between either cage partner and animals in other cages. ANOVA for repeated measures on the segment 0.4-0.9 s was significant (F(1,19)=11.36, p=0.003).

**Figure 4.**
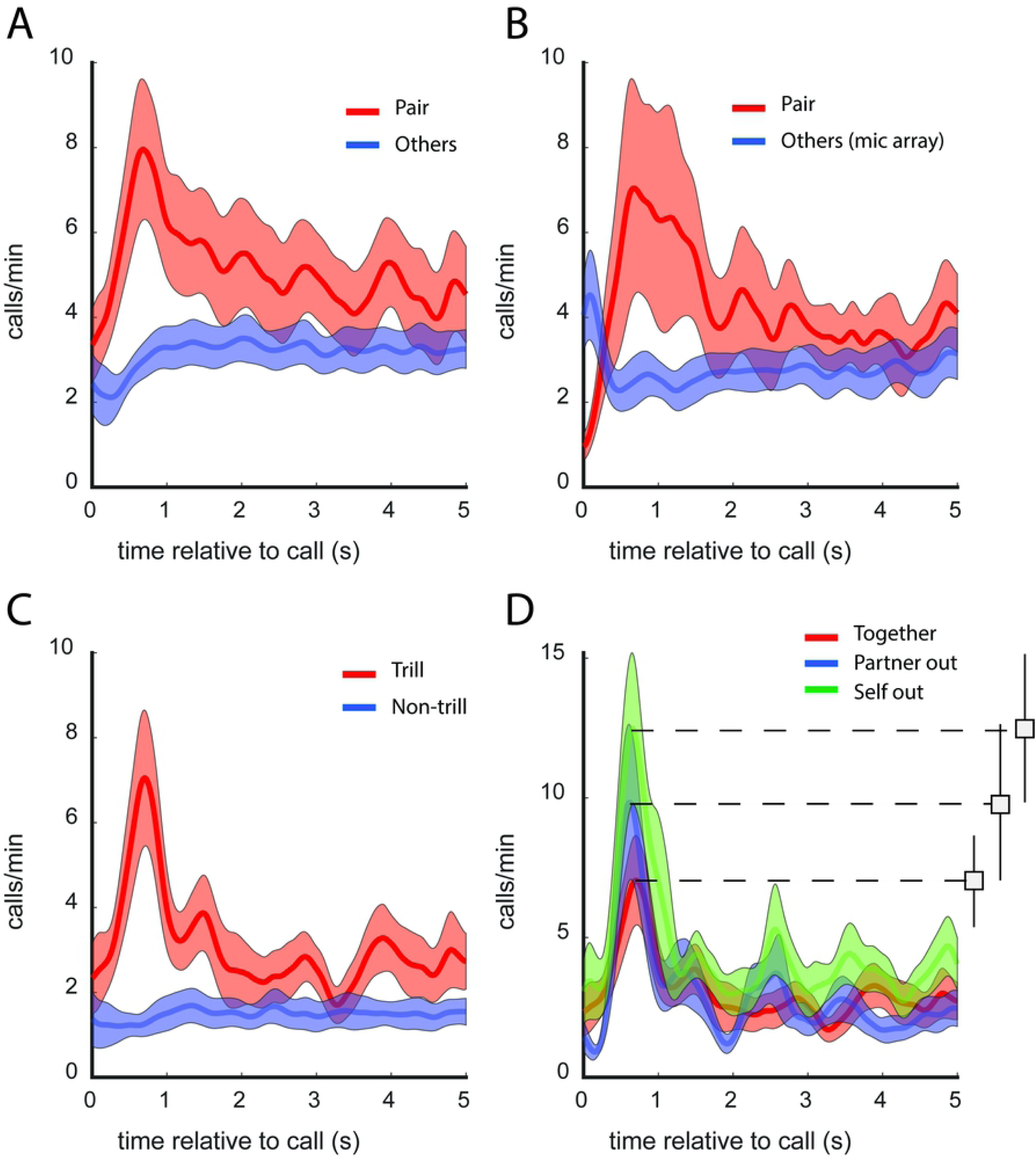
Timing of calls relative to calls from partner and other animals. **A**. Call rate in relation to calls from the partner and other animals in the room. There is a transient increase in call rate shortly after calls from the partner, but not after calls from other animals. Shaded areas indicate the standard error of mean. **B**. Results from a subset of 6 animals (3 pairs) with an array of microphones in the room, showing a similar pattern as in A, confirming the result found with wearable recorders alone. **C**. Trills versus all other call types combined. The peak is absent for other call types, showing that the temporal relation at this latency mainly involves trill calls. **D**. Trill call rate in relation to calls from the partner when together in the home cage, the partner is out, or the animal him/herself is out. This shows that responsiveness to the partner’s calls is strongest when the animal him/herself is out.

To confirm that the separation between calls from the pair and calls from other animals using the wearable recorders was correct, a subset of recordings was done with additional microphones in the room. A time-difference-of-arrival method was employed to separate between the cage under study and other cages. There was 87% agreement between attributions from this method and the wearable recorders. Figure 4B shows call rate contingent upon the partner as obtained from the wearable recorders, and call rate contingent upon others as obtained from the microphone array. This shows a similar pattern to our observation from only the voice recorders: A peak in call interaction for cage partners but not between either cage partner and other animals (Wilcoxon signed rank test on segment 0.4-0.9 s: Z=-2.2, P=0.028). The peak seen at 0 s for ‘others’ is because some calls that were assigned to ‘others’ by the mic array were assigned to the partners by the wearables.

Next, we split the data into different call types (Figure 4C). When using only trill calls, there is a peak, similar to when using all calls, but when using non-trill calls, there is no peak (Rep Measures ANOVA using mean 0.4-0.9 s, F(1,19)=11.6, p=0.003). Therefore, in these circumstances the temporal relationship involves trill calls rather than other call types. There was not enough data to split non-trill calls into different call types for this analysis. The majority of animals (19 out of 20) had a significant peak in trill rate following trills from the partner (5 of 5 females, 14 of 15 males). The mean latency of the peak was 629 ms after the partner’s call.

To examine whether responsiveness to the partner changes when animals are separated, we compared the peaks in call rate between the separation conditions (together, partner out, self out). As Figure 4D shows, the peak was highest in the ‘self out’ condition, indicating that animals were most responsive to calls from their partner when they themselves were outside the home cage. ANOVA for repeated measures confirmed the main effect of separation condition (F(2,38)=4.67, p=0.015).

Next, we analyzed spectral features of the trill calls. Figure 5A shows the spectrogram, spectrum and the fundamental frequency F0 of a single trill call. Mean spectrograms for the population are shown in Figure 5B. Separation had an effect on F0 (Repeated measures ANOVA F(2, 38) = 7.05, p =0.002.). The median F0 for *Self out* (7946 Hz) was 969 Hz higher than for *Together* (6977 Hz). For intuitive reference, this is approximately one whole tone higher in musical terms (23). There was no significant change in vocalization intensity.

**Figure 5.**
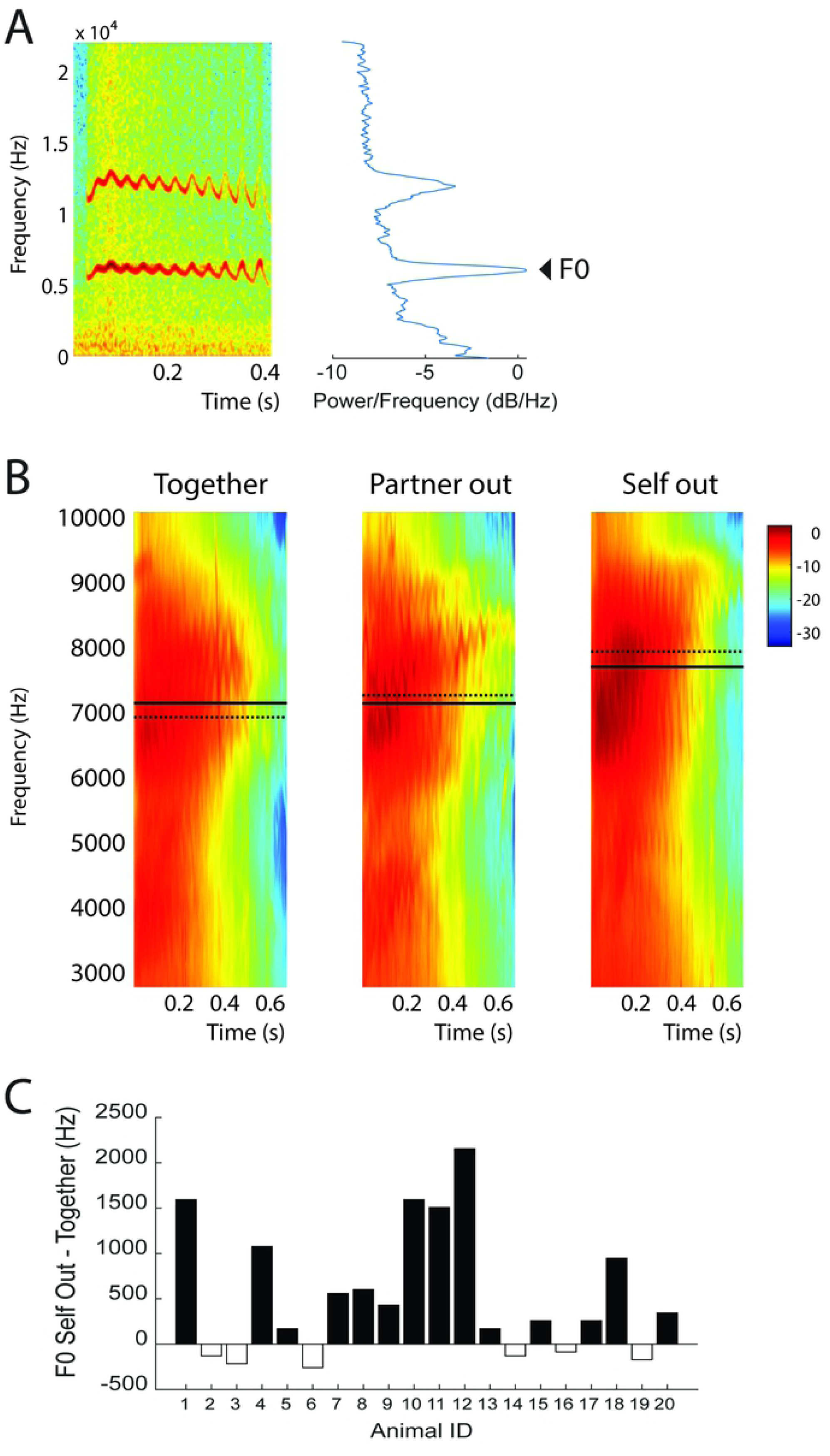
Spectral analysis **A** Example spectrogram (left) and spectrum (right) of a single trill call. The fundamental frequency F0 is indicated. Trill calls often have multiple harmonics. The fundamental frequency is the first (and lowest) harmonic. **B**. Population mean spectrograms of trill calls in the 3 conditions. The solid and dashed horizontal lines indicate the mean and median F0. **C**. F0 for the ‘self-out’ condition minus the ‘together’ condition for each animal.

## Discussion

We have shown that we can monitor close range vocal interactions in the home cage in freely moving, socially housed marmosets using wearable voice recorders. In these circumstances, trill calls were by far the most common call type. Trill call interactions are thought to serve as a means to signal one’s presence (6,7). Therefore, we expected that being separated while still at close range would affect trill calls. For each animal, we compared a condition in which they were together with their partner, a condition where the partner was in a separate cage, and a condition where the animal itself was in a separate cage. Separation resulted in an increase in trill call rate, while other call types were unaffected. When the animal him/herself was in the separate cage, trill call rate increased the most.

There was a temporal relationship between vocalizations from the members of each pair as evidenced by a peak in call rate just after calls from the partner. No such relationship was observed between either member of the pair and other animals in the room. It is safe to assume animals have a stronger social bond with their cage partner than with animals in other cages. Therefore, the strength of the vocal interactions we observe might reflect the strength of the social bond. In pygmy marmosets (7,11,16), macaques (24,25) and squirrel monkey (26,27), contact calls have been shown to be affiliative and correlated with social bonds. The present data does not allow attribution of vocalizations from other animals in the room to specific individuals. Thus, although there was no temporal coordination of calls between the cage partners under study and the other animals in the room as a group, it is possible that there were interactions between either cage partner and specific individuals in other cages. More research is necessary to test that possibility.

The call exchanges we observed predominantly involved trill calls. The estimated latency of 629ms for trill responses to trills from the partner is in agreement with previous work (6). When the animal was in the separate cage, the height of the peak for that animal increased, indicating that separation makes the animals more responsive to calls from their partner. This makes sense if the function of trill call interactions is to signal one’s presence. In the wild, a contact call system in which a response is expected at a certain time interval may serve to determine whether an animal is missing (16).

We find that the fundamental frequency, F0, of trills increases when animals are placed outside the cage. Increased arousal has been linked to increases in F0 in many species including cattle, pig, cat, hyena, seal, dolphin, bat, macaque, baboon, squirrel monkey, guinea pig, marmot and tree shrew (17). In humans, F0 is among the features that correlate with cognitive workload (28), but also with anger, fear and joy, while sadness is more associated with a decrease in F0 (18). In marmosets, Schrader and Todt showed that increasing levels of isolation resulted in increases in the F0 of phee calls (29). The fact that F0 was only increased when animals were themselves outside the home cage and not when their partner was out, indicates that F0 increase is not merely due to an increase in distance between the animals. It is possible that increased arousal results in increased muscle activity in the larynx, thus increasing the fundamental frequency (30). If modulations in audio features vary systematically with arousal or emotional state, they may also have communicative value. Humans can detect emotions from vocal cues including pitch (31,32). When normal pitch modulation (prosody) is reduced, as is often the case in autism (33), Parkinson’s disease (34), and schizophrenia (35), speech becomes monotonic, which negatively affects verbal communication. Although the current study shows that social context affects timing and pitch, further research is needed to determine whether aberrant timing and pitch negatively affect social interactions.

## Acknowledgement

The authors wish to thank the following for their generous support: The Stanley Center for Psychiatric Research, the Poitras Center for Psychiatric Disorders Research, the Simons Center for the Social Brain, the Tan-Yang Center for Autism Research, and NIH (Awards No. 1R01MH111916-01A1 and 2P50MH094271-06 to G.F. and 5R01MH111916-02 to R. D.).

